# *In vitro* biofilm formation only partially predicts beneficial *Pseudomonas fluorescens* protection against rhizosphere pathogens

**DOI:** 10.1101/2024.12.17.628960

**Authors:** Yang Liu, Alexandra D. Gates, Zhexian Liu, Quinn Duque, Melissa Y. Chen, Corri D. Hamilton, George A. O’Toole, Cara H. Haney

## Abstract

Plant roots form associations with both beneficial and pathogenic soil microorganisms. While members of the rhizosphere microbiome can protect against pathogens, the mechanisms are poorly understood. We hypothesized that the ability to form a robust biofilm on the root surface is necessary for the exclusion of pathogens; however, it is not known if the same biofilm formation components required *in vitro* are necessary *in vivo. Pseudomonas fluorescens* WCS365 is a beneficial strain that is phylogenetically closely related to an opportunistic pathogen *P. fluorescens* N2C3 and confers robust protection against *P. fluorescens* N2C3 in the rhizosphere. We used this plant-mutualist-pathogen model to screen collections of *P. fluorescens* WCS365 increased <u>a</u>ttachment <u>m</u>utants (*iam*) and <u>s</u>urface <u>a</u>ttachment <u>d</u>efective (*sad*) transposon insertion mutants that form increased or decreased levels of biofilm on an abiotic surface, respectively. We found that while the *P. fluorescens* WCS365 mutants had altered biofilm formation *in vitro*, only a subset of these mutants, including those involved in large adhesion protein (Lap) biosynthesis, flagellin biosynthesis and O-antigen biosynthesis, lost protection against *P. fluorescens* N2C3. We found that the inability of *P. fluorescens* WCS365 mutants to grow *in planta*, and the inability to suppress pathogen growth, both partially contributed to loss of plant protection. We did not find a correlation between the extent of biofilm formed *in vitro* and pathogen protection *in planta* indicating that biofilm formation on abiotic surfaces may not fully predict pathogen exclusion *in planta*. Collectively, our work provides insights into mechanisms of biofilm formation and host colonization that shape the outcomes of host-microbe-pathogen interactions.

## Introduction

Microbiota play a key role in plant and animal defense against pathogens, both by modulating the immune system of their host and by excluding pathogens [1, 2]. Pathogen exclusion can occur through antagonism and the production of antimicrobials, or through niche competition and exclusion [3]. Niche exclusion can occur through direct physical competition, for instance by occupying space, or alternatively, through more efficient use of nutrients. While many antimicrobials made by microbiota that target pathogens have been identified, how microbiota exclude pathogens is poorly understood.

Biofilm formation has been implicated in both microbial virulence, as well as microbiota-mediated exclusion of pathogens [4]. Biofilms are comprised of mechanistically diverse extracellular matrices consisting of proteins and exopolysaccharides that are formed by microbes for biotic and abiotic surface attachment [5, 6]. For plant-associated microbiota, biofilm formation is required for rhizosphere colonization. For instance, a reverse genetics screen of *Bacillus* biofilm determinants identified that many *in vitro* biofilm components are also required to colonize plants [7]. Similarly, a forward genetic screen in a beneficial *Pseudomonas* strain found the large adhesion protein LapA is required to colonize corn roots [8]. LapA is also required for biofilm formation *in vitro* [9] indicating that there might be overlapping mechanisms between biofilm formation and host association. Biofilm formation by microbiota is also associated with the prevention of fungal pathogen invasion of amphibians [10]. However, while biofilms have been extensively studied *in vitro*, there is limited data as to whether biofilm formation and protection against plant pathogens *in vivo* require the same mechanisms.

Reductionist model plant-microbiota-pathogen systems have facilitated the identification of mechanisms by which microbiota can protect hosts from pathogens [11]. A previously described model system consisting of a beneficial *Pseudomonas fluorescens* strain WCS365, a closely related pathogen *Pseudomonas fluorescens* N2C3, and the model plant *Arabidopsis thaliana* (Arabidopsis) was used to identify mechanisms required by mutualists for protection against pathogens [12]. For example, plant colonization through a two-component system ColR/S and LPS core polysaccharide modification was shown to be required for WCS365-mediated protection against N2C3 [12]. We hypothesized that robust biofilm formation by the beneficial strain *P. fluorescens* WCS365 would be required for protection against pathogenic N2C3.

To identify bacterial biofilm components necessary for pathogen protection, we screened two previously described, but only partially characterized, collections of *P. fluorescens* WCS365 biofilm transposon insertion mutants for pathogen protection [9, 13]. These include mutants with decreased (*sad,* <u>s</u>urface <u>a</u>ttachment <u>d</u>efective mutants) and increased (*iam,* increased <u>a</u>ttachment <u>m</u>utants) biofilm formation on abiotic (plastic and glass) surfaces [14]. From this screen, mutations in genes encoding the large adhesion protein A (LapA) system were described as promoting biofilm formation by *P. fluorescens* [14]. Because LapA was also previously implicated in plant association [8], and because the majority of mutants in the *sad* and *iam* collections have had limited characterization, we hypothesized that this library is a source of novel rhizosphere colonization determinants required to exclude pathogens. Furthermore, the *iam* mutants provide the opportunity to determine if increased biofilm formation can enhance pathogen protection, or rather, will result in colonization defects because of mis-regulation of biofilm formation [15].

By mapping the genetic location of transposon insertions in the *P. fluorescens* WCS365 *iam* and *sad* biofilm libraries, we identified mutations in both previously described and novel biofilm formation components. While the *iam* and *sad* mutants had altered biofilm formation *in vitro*, we found only a subset of these mutants lost the ability to protect against pathogens *in planta*.

These results suggest that only a subset of biofilm components required *in vitro* are required for plant-protective functions *in planta,* and that *in vitro* biofilm formation and *in planta* pathogen protection use only partially overlapping mechanisms.

## Results

### Rescreening a collection of *P. fluorescens* biofilm mutants identified genes required for pathogen-protection

We set out to determine if *P. fluorescens* WCS365 mutants with increased or decreased biofilm formation [9] could still protect plants from the closely related *P. fluorescens* pathogen N2C3 [12]. *In vivo* biofilm formation is challenging to quantify directly, and we reasoned that protection against pathogens would provide a readout for the symbiotic ability of the *P. fluorescens* WCS365 biofilm mutants. As not all of the *P. fluorescens* WCS365 *iam* and *sad* library transposon insertions sites were mapped previously, we first mapped the transposon insertions by arbitrary primed PCR [14]. The PCR products were sequenced and aligned to the *P. fluorescens* WCS365 reference genome to determine the location of the transposon insertions [16]. Among a total of 62 mutants, we identified insertions in 35 unique genes, the majority of which were previously implicated in biofilm formation (Table S1, Figure 1A). The majority of insertions were within 4 genetic loci including 10 insertions within a predicted O-antigen biosynthesis operon (*wbpADE/wzt/wphH/wapH/wbpJ*; [17][18]), 6 insertions in a flagellin biosynthesis operon (*fliK/fliN/fliP/fliR*; [19][20], 16 insertions in large adhesion protein biosynthesis locus (*lapDAEB;* [16, 21]), and 7 insertions in an LPS biosynthesis operon (RS13585/*warR;* [22]) (Figure 1A). Additional genes with multiple insertions from the screen include *pvdQ* [23]*, hisD,* and *rlmN* (Table S1). There were 7 insertions in genes and operons that only had a single insertion including in intergenic regions or hypothetical proteins (Table S1). As we had less confidence that these insertions were responsible for the observed phenotype, these singletons were not considered further except for *clpP* [14, 24] which has been previously characterized in WCS365 biofilm formation. Collectively these findings indicate that the *iam* and *sad* library robustly identified known biofilm components in *P. fluorescens* and related bacterial taxa.

**Figure 1.**
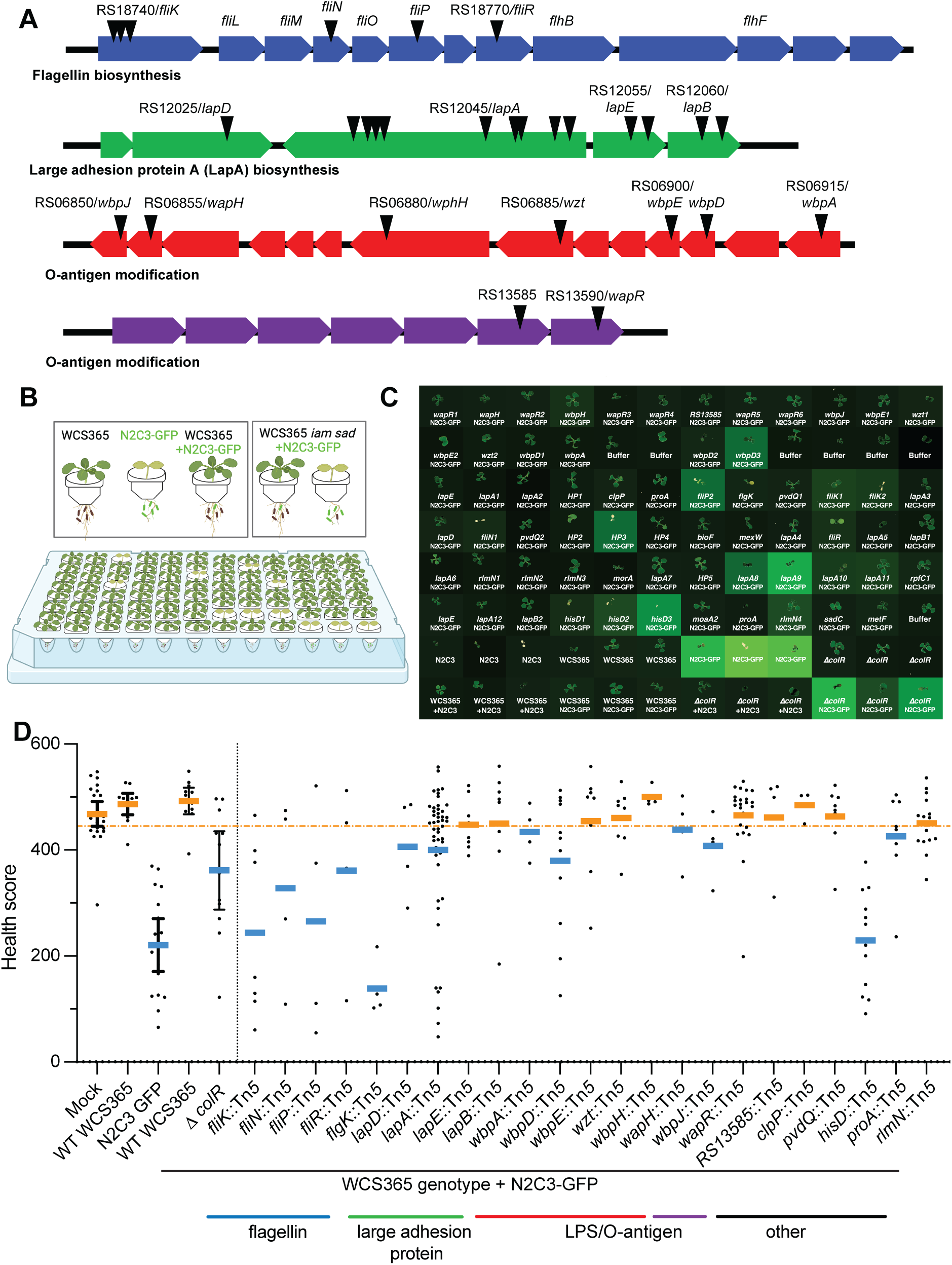
A high throughput screen identified *P. fluorescens* WCS365 biofilm mutants that cannot protect plants from a pathogen. A) Schematic of four major operons identified and characterized in the screen including flagellin biosynthesis, large adhesion protein, and 2 operons involved in O-antigen modification. Triangles indicate approximate locations of transposon insertions. B) Schematic of high-throughput pathogen protection screen where plants are grown in 96-well plates and treated with either a pathogen, a commensal, or in combination. Increased attachment mutants (*iam*) and surface attachment defective (*sad*) mutants of WCS365 were screened using a 96-well plant health quantification assay. Plants were inoculated with a 1:1 ratio of WCS365 *iam* or *sad* libraries and the N2C3 pathogen expressing GFP from a plasmid. C) Plant health and bacterial fluorescence was quantified. Plant health was quantified by scanning plants with a high-resolution scanner and quantifying plant size and color. Growth of the bacterial pathogen was quantified by reading GFP fluorescence. D) Plant health was quantified as a function of how large and how green plants are at the end of the assay period. Using this metric, wildtype WCS365 results in largely healthy plants even with the N2C3 pathogen is present. A previously described WCS365 *ΔcolR* mutant that cannot protect plants was used as a control; the dotted orange line indicates the threshold to detect *ΔcolR* mutants with 95% confidence, which was used as a cutoff in this preliminary screen. Bars are colored by whether strains were protective (orange) or non-protective (blue). Each dot represents a single plant from 4 independent replicates, and genes with multiple insertions are pooled. Bars indicate the mean, and then error bars represent the 95% confidence intervals of the control.

### Correlation between *in vitro* biofilm formation and plant protection

We hypothesized that biofilm formation would be positively correlated with plant colonization and protection against pathogens. However, we also hypothesized that increased biofilm formation, could potentially be detrimental for plant colonization and pathogen protection. To screen the *P. fluorescens* WCS365 *iam* and *sad* mutants for protection against the pathogenic *P. fluorescens* N2C3, we made use of a high-throughput plant protection assay [25]. In this assay, WCS365 protects against N2C3 and results in healthier plants and low levels of N2C3 abundance; both bacterial abundance and plant health can be readily quantified (Figure 1B). We inoculated plants with wildtype WCS365 or individual *iam* or *sad* mutants in combination with *P. fluorescens* N2C3 containing a plasmid expressing GFP, and performed the assay with 4 biological replicates (Figure 1B). We then quantified plant health and N2C3-GFP fluorescence to determine if the WCS365 mutants were no longer able to protect the plant and if N2C3 was able to grow in the rhizosphere (Figure 1B and 1C).

We found that a subset of *iam* and *sad* mutants could no longer protect plants against disease caused by *P. fluorescens* N2C3 as measured by a decreased plant health score (Figure 1C,D). Interestingly, nonprotective *iam* and *sad* mutants included both those with increased and decreased biofilm formation. Nonprotective *sad* mutants included insertions in genes involved in large adhesion protein biosynthesis (*lapDAEB)*, consistent with previous descriptions of a role for the Lap adhesion system in plant colonization [7] and flagellar biosynthesis (*fliKNPR* and *flgK*). Nonprotective *iam* mutants included those with insertions in lipopolysaccharide (LPS) modification (*wbpAD*). Conversely, a number of other mutants including those involved in LPS modification retained protective ability. Collectively, these findings indicate that some, but not all, genes required for *in vitro* biofilm formation are required for pathogen protection *in planta*.

### Decreased *in planta* fitness partially explains loss of protection against pathogens

In our high throughput plant protection assay, each well contains plant root exudates that can support the growth of bacteria. Under these conditions, wildtype *P. fluorescens* WCS365 outcompetes pathogenic N2C3 in the rhizosphere resulting in WCS365 dominating the final community in each well [12, 25]. Here, co-inoculation with WCS365 resulted in low growth of N2C3, and N2C3-GFP fluorescence was not detectable above the background detection limit (Figure 2A). We hypothesized that nonprotective *iam* and *sad* mutants would be associated with increased growth of the N2C3 pathogen, but potentially not a change in growth of the WCS365 mutant.

**Figure 2.**
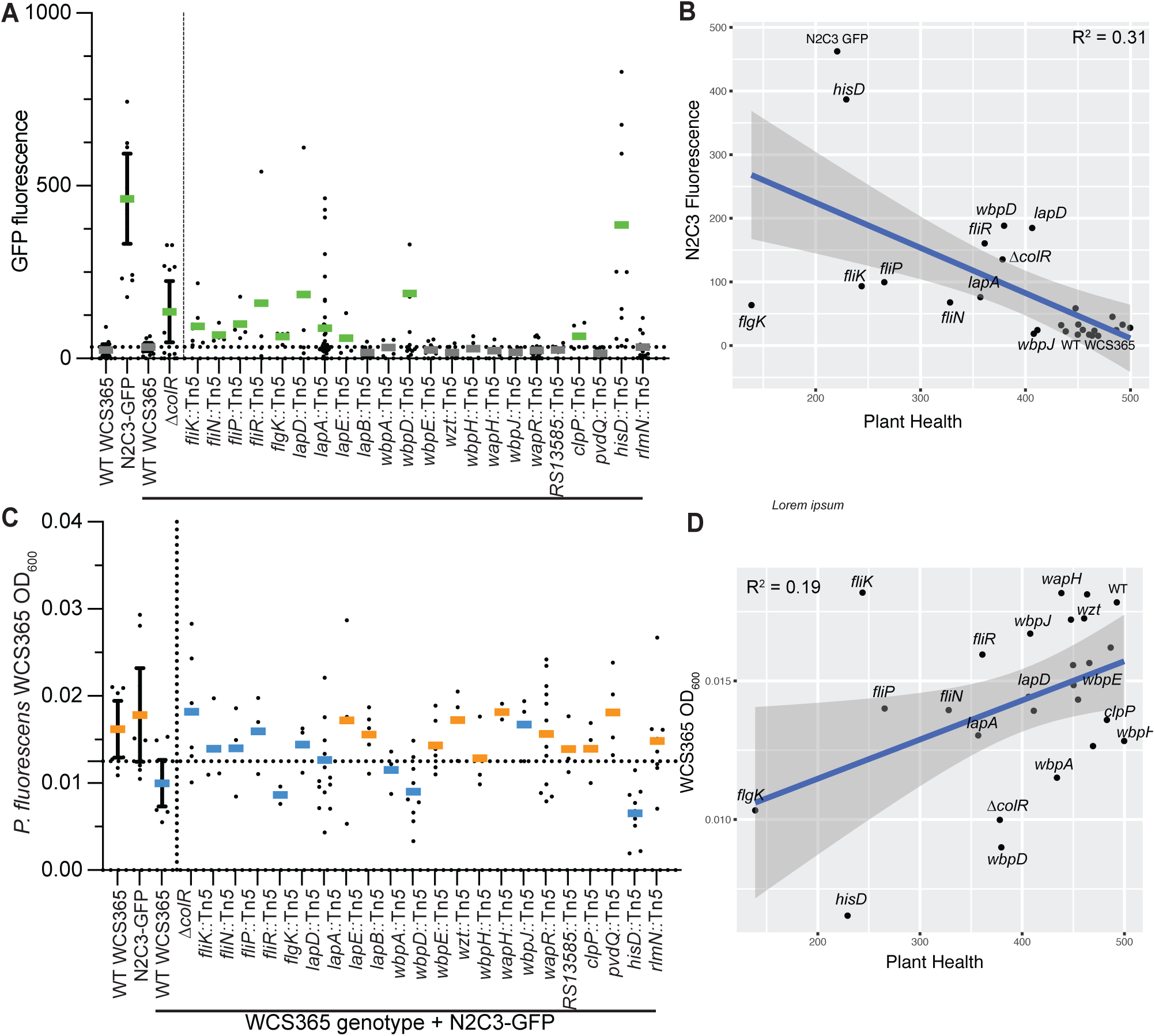
Decrease in commensal fitness and increase in pathogen growth in planta contribute to loss of protection. A) Quantification of N2C3-GFP signal in competition with *P. fluorescens*. The horizontal dashed line indicates the background detection limit; those with signal above background are shown in green. B) Increased growth of the N2C3-GFP strain partially explains a decrease in plant health. C) Growth of *P. fluorescens* WCS365 *iam* and *sad* mutants in competition with N2C3. The dashed line shows the 95% confidence interval of wildtype WCS365 growth. Bars are colored by whether strains were protective (orange) or non-protective (blue) from Figure 1. D) Decreased growth *in planta* explains some but not all of the loss of protection.

We first measured pathogen growth by quantifying N2C3-GFP signal in each well. We found that several WCS365 *iam* or *sad* mutants that resulted in decreased health scores when competed against N2C3 also had increased N2C3-GFP fluorescence (Figure 1C and 2A) suggesting that the ability to limit pathogen growth may play an important role in mutualist-mediated plant health. To test whether all WCS365 mutants that lost protection also had corresponding increases in N2C3 abundance, we performed a linear regression between N2C3-GFP fluorescence and plant health. We found a significant negative correlation between N2C3 abundance as measured by GFP fluorescence signal and plant health (R^2^= 0.31, *p* = 0.0035; Figure 2B). Interestingly, insertions in the *hisD* auxotrophic mutants resulted in a similar level of N2C3-GFP fluorescence as N2C3-GFP alone suggesting these mutants likely had low growth in the rhizosphere (Figure 2A). In contrast, the majority of flagellin mutants failed to protect plants, but without a corresponding increase in GFP signal (Figure 2B). This suggests that the flagellin mutants may still inhibit N2C3 growth, but that loss of flagellar motility may result in a loss of pathogen protection due to decreased plant colonization. Altogether, these results are consistent with loss of protection through diverse mechanisms.

Increase in N2C3 growth and decrease in plant health in the presence of *iam* and *sad* mutants could be a result of rhizosphere fitness defects or due to loss of niche occupation because of altered biofilm formation*. In planta,* we found that a subset of the nonprotective strains had reduced rhizosphere growth consistent with rhizosphere fitness defects (Figure 2C). We found a positive correlation between plant health and growth of WCS365, although less robust than the correlation between N2C3 abundance and plant health (R^2^ = 0.19, *p* = 0.028; Figure 2D). This suggests that some *iam* or *sad* mutants, such as the *hisD* mutants, were unable to grow *in planta*, which explains their inability to exclude the N2C3 pathogen. Others, such as a strain carrying a mutation in the *fliK* gene, still grew to fairly high levels but cannot protect plants against N2C3. Interestingly, similar to the previously described *ΔcolR* mutant, the *wbpAD* mutants, which are involved in O-antigen biosynthesis, grow to an intermediate level in the rhizosphere. ColR regulates genes involved in LPS modification, although not specifically *wbpAD* [26]. The co-clustering of these mutants for rhizosphere fitness and plant health suggests that *wbpAD* and *colR* genes may contribute to rhizosphere fitness through related processes.

To determine if growth defects of *iam* or *sad* mutants were rhizosphere specific, or the mutants have generalized growth defects, we performed *in vitro* growth curves with these mutants in LB medium. We found that the majority of mutants grew to similar levels as wildtype bacteria (Figure S1) and that those with significant, although modest changes in growth did not correlate with those that could no longer protect (R^2^ = 0.0024, *p* = 0.82; Figure S1). Collectively, these findings indicate that both the ability to exclude pathogens as well as rhizosphere fitness contribute to WCS365-mediated protection.

### Some biofilm mechanisms are conditionally required for protection against pathogens

We previously found that there is not a perfect correlation between genes required for pathogen protection in hydroponics compared to a solid surface assay [12, 26]. While the hydroponic assay has the benefit of being high throughput, it may not recapitulate all aspects of growing *in planta*, including the necessity to move across liquid-air interface and solid surfaces. As a result, we repeated protection assays for 19 mutants representing the majority of genes and operons identified in the screen using a solid surface plant assay [12]. In these assays, plant fresh weight is a read-out for protection. Our results show N2C3 significantly stunts plant growth, while plants treated with WCS365 or a 1:1 ratio of WCS365 and N2C3, have similar fresh weights as buffer-treated plants (Figure 3A). To test for pathogen protection, we co-inoculated *P. fluorescens* WCS365 or WCS365 mutants with *P. fluorescens* N2C3 in a ratio of 1:1 along the plant roots.

**Figure 3.**
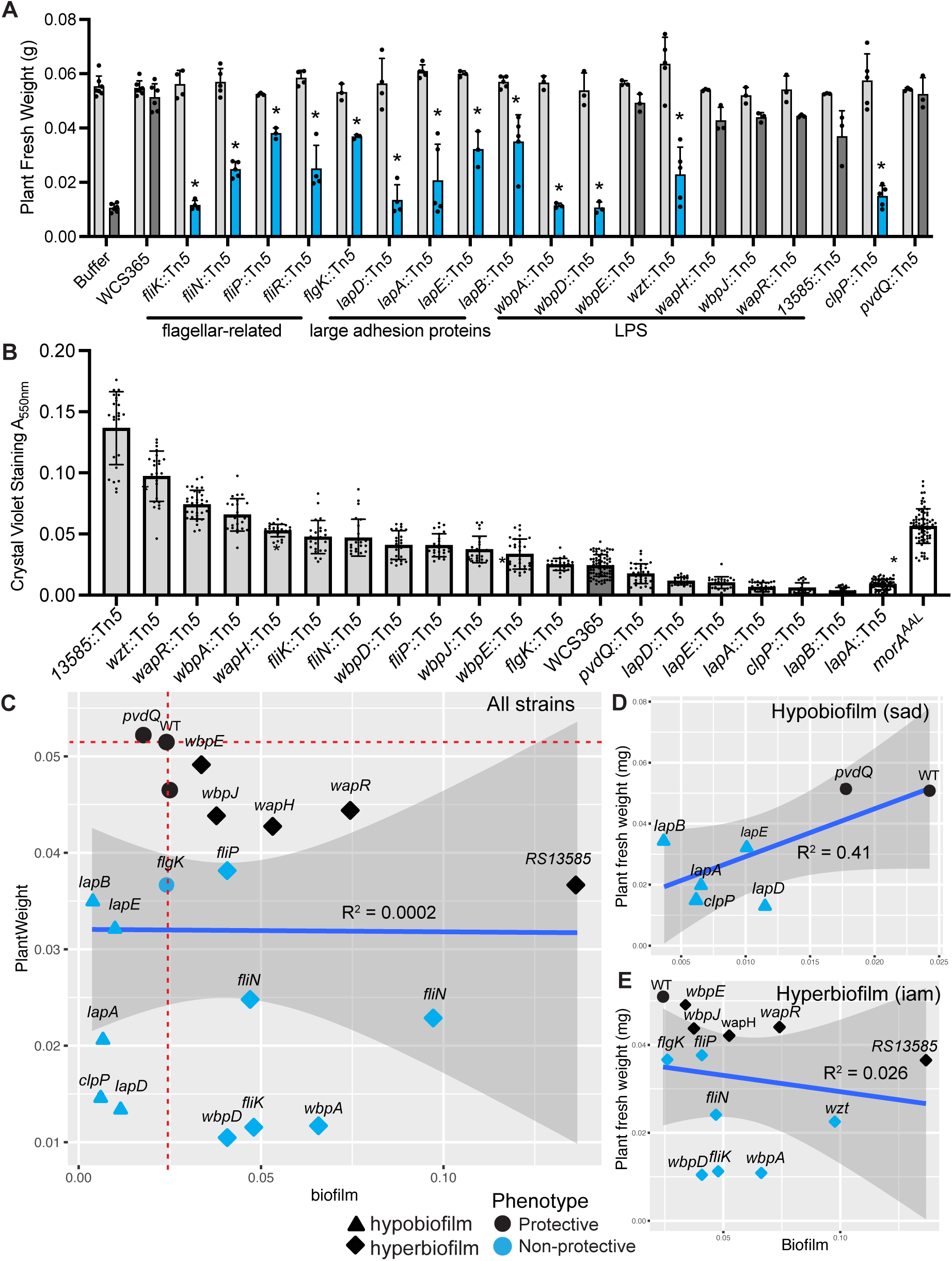
The extent of *in vitro* biofilm biomass does not predict protection against a pathogen *in planta.* **A)** From among the 65 *P. fluorescens* WCS365 biofilm formation mutants in the *iam* and *sad* mutant collections (Table 1), 21 mutants were re-tested for their ability to protect the rhizosphere against disease caused by the *P. fluorescens* N2C3 pathogen. In a competition assay using *P. fluorescens* WCS365 mutants with *P. fluorescens* N2C3 in a ratio of 1:1, 12 mutants lost protection in the rhizosphere. Light grey indicates buffer (MgSO_4_) or *P. fluorescens* WCS365 wildtype bacteria or mutants in mono-association, and blue or dark grey indicates *P. fluorescens* N2C3 or biofilm mutants in competition against *P. fluorescens* N2C3. Each dot represents an average of three technical replicates as one independent biological replicate. The assay was repeated at least three times. Statistical significance was determined by two-way ANOVA comparing different strains with N2C3 treatment, *p* < 0.001 = *. Error bars show standard deviation. B) *in vitro* biofilm biomass was quantified using a crystal violet assay. The *sad-51/lapA::*Tn*5* and *morA*^AAL^ mutations are previously described hypo- and hyper-biofilm controls, respectively. Each dot represents a technical replicate from one independent biological replicate and each biological replicate includes 8 technical replicates. The assay was repeated at least three times. Statistical significance was determined by one-way ANOVA comparing different strains with WCS365 treatment followed by a Tukey’s HSD test \**p* < 0.05. All error bars show standard deviation. C) Correlation between *in vitro* biofilm formation versus plant weight. The plant weight was used as a readout for protection and whether the tested mutant can outcompete *P. fluorescens* N2C3 in the rhizosphere or not. Each dot represents the average of at least three independent biological replicates quantifying biofilm formation and plant weight. D-E) Independent linear regressions were performed for just the *sad* (D) or *iam* (E) mutants. Linear regression analysis was performed in R by ggplot2. Only a positive correlation between the *in vitro* hypobiofilm formation and protection was found.

We found that 13 of the 19 *P. fluorescens* WCS365 mutants lost the ability to protect against *P. fluorescens* N2C3 as indicated by a significant decrease in fresh weight relative to the protective wildtype *P. fluorescens* WCS365. These include mutants with insertions in genes coding for the Lap system (*lapDAEB*) [27], motility (*fliK*, *fliR* and *fliN*), lipopolysaccharide (LPS) modifications (*wbpAD*, *wzt*)[28, 29] and a protease encoded by the *clpP* gene [14] (Figure 3A). The remaining *P. fluorescens* WCS365 mutants maintained pathogen protection as indicated by no significant difference in plant weight when co-inoculated *P. fluorescens* N2C3 (Figure 3A). While all *iam* and *sad* mutants defective in plant protection against N2C3 in hydroponics were also required in a solid surface plant assay, several additional genes, when mutated, were shown to be conditionally required on the solid surface plant assay including *clpP, wzt,* and *lapEB.* These results indicate a potential differential requirement for some components of biofilm formation involved in pathogen protection on plants growing on a solid surface. As plant growth on solid agar may more closely mimic soil where microbes must navigate air-liquid interfaces and movement along soil particles, the solid surface plant assay may more closely mimic soil conditions and reveal the importance of additional biofilm genes in pathogen protection.

### There is no correlation between the extent of biofilm formed *in vitro* and pathogen protection *in planta*

We found that a subset of *P. fluorescens* WCS365 mutants from both the *iam* and *sad* libraries lost protection against *P. fluorescens* N2C3, and therefore there did not appear to be a correlation between the extent of biofilm biomass produced and protection. Using the same subset of 19 mutants retested on the solid plate assay, we robustly quantified *in vitro* biofilm formation and *in planta* competition.

Using a modified crystal violet assay [30] we validated the previously described increased or decreased biofilm phenotypes for the 19 *P. fluorescens* WCS365 mutants *in vitro*. The assay conditions were nearly identical to the original assay including the same media, but the biofilm was allowed to proceed overnight for 18 hours, instead of 10 hours [9], to reflect the longer interaction time with a plant. We found 13 mutants, including components of flagellin biosynthesis and LPS modification, formed significantly higher amounts of biofilm than wildtype *P. fluorescens* WCS365 (Figure 3B). We found 6 mutants, primarily in the Lap system, formed lower biofilm amounts (Figure 3B). It is noteworthy that in the repeat of the biofilm assays the flagellin biosynthesis mutants formed increased biofilm, while they were originally described as reduced for biofilm formation [9]. Quantitative differences in biofilm formation have previously been described for *Pseudomonas* flagellin mutants as a function of time and media [31]. As the replicate assay was performed in the same media as the initial assay but over 18 hours instead of 10 hours as in the initial screen, we hypothesize the longer incubation time is the most likely explanation for the difference in mutant phenotype. The remaining mutant phenotypes are consistent with descriptions of biofilm phenotypes from the *iam* and *sad* mutants and previous descriptions of biofilm components [9].

To determine if the extent of the *in vitro* biofilm biomass formed could predict *in planta* pathogen protection, we performed a linear regression between the plant weight, as a readout for protection, and *in vitro* biofilm formation. No significant correlation between the two phenotypes was found, suggesting that the extent of the biofilm formed by a strain *in vitro* does not directly predict its protective ability *in planta* (R^2^ = 0.0002, *p* = 0.98; Figure 3C). As non-protective WCS365 mutants included both decreased (*sad*) and increased (*iam*) attachment mutants, we formed separate regression analyses to test if there was a correlation between mutants that formed decreased or increased biofilm. Interestingly, we found a trend between decreased biofilm formation and decreased protection (R^2^=0.5, *p* = 0.77; Figure 3D) suggesting that loss of biofilm formation results in inability to protect plants from N2C3. In contrast, there was no correlation between increased WCS365 biofilm formation and protection against N2C3 (R^2^ = 0.046, *p* = 0.46; Figure 3E). These results are consistent with diverse mechanisms of biofilm formation *in vitro*, that may not precisely correlate with functions *in planta*.

### Biofilm components are required for protection against diverse rhizosphere pathogens

*P. fluorescens* N2C3 is an opportunistic pathogen of plants and produces a lipopeptide toxin [32]. Previously described *P. fluorescens* WCS365 genes required for protection against N2C3 are also required for protection against a virulent rice pathogen *Pseudomonas fuscovaginae* SE-1, which uses a similar toxin-based virulence mechanism to cause disease [12]. *Pseudomonas aeruginosa* is an opportunistic pathogen of both plants and animals but uses distinct virulence mechanisms from N2C3 and SE-1 [33]. We tested whether the same genes required to protect against N2C3 were also required to protect against *P. fuscovaginae* SE-1 and *P. aeruginosa* PAO1 by choosing 3 mutants affecting diverse processes, including *lapA::*Tn5 (large adhesion protein, hypobiofilm), *wzt*::Tn5 (LPS modification, hyperbiofilm) and *clpP*::Tn5 (protein turnover, hypobiofilm). We found that wildtype WCS365 robustly protects from both plant biomass decreases (Figure 4A) and root stunting (Figure 4B-C) by all three pathogens. We found consistent loss of protection by the *lapA::*Tn5, and *wzt::*Tn5 and *clpP::*Tn5 mutants against both N2C3 and SE-1. Interestingly, none of the genes were required for protection against PAO1 as measured by no significant reduction in plant biomass relative to the mutant alone. The only exception was that the *lapA* mutant, which did not fully protect against PAO1-mediated root stunting. Interestingly, we found that *clpP* mutant itself resulted in plant root stunting, however the fresh weight of plants inoculated with the *clpP* mutant did not differ from buffer treated plants (Figure 4). Collectively these findings indicate that precise regulation of biofilm formation is required for commensals to colonize plant roots and protect plants against pathogens of agronomic importance.

## Discussion

Here, we screened a collection of *P. fluorescens* WCS365 transposon insertion mutants with altered biofilm formation on abiotic surfaces using a previously described plant-commensal-pathogen model [34, 35]. By employing a high-throughput assay that measures fluorescence and absorbance, we were able to monitor the plant health as well as the individual growth of both a pathogen and a commensal coexisting in the rhizosphere [25]. We found that not all the mutants in the *iam* and *sad* libraries lost protection of plants against a pathogen, suggesting that not all biofilm formation mechanisms *in vitro* are required for protection against pathogens *in planta*. We found that the plant health is more strongly correlated with the abundance of the pathogen than the commensal, as some biofilm mutants were able to maintain their growth but failed to provide protection. Our results suggest that protection requires close association of the protective microbe with the plant and not just growth in the rhizosphere.

**Figure 4.**
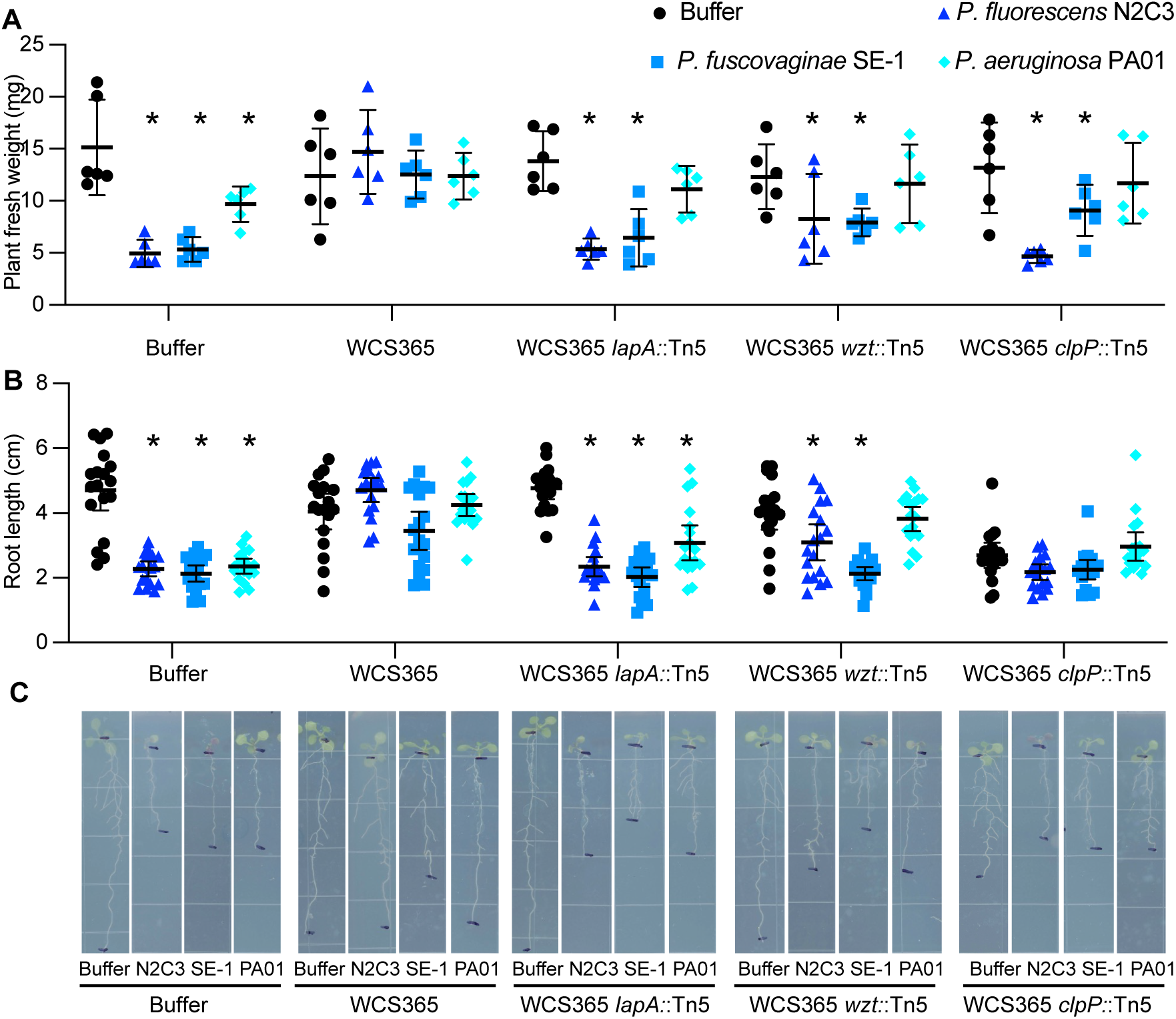
Biofilm mutants have protection defects against diverse pathogens. To determine if mechanisms of *P. fluorescens* WCS365 excluding pathogens are also important for *Pseudomonas* pathogens with distinct virulence mechanisms, we tested mutants in three distinct processes for their ability to protect against *Pseudomonas fuscovaginae* SE-1 and *Pseudomonas aeruginosa* PAO1. Plant biomass (A) and root length (B) were quantified with representative images shown in panel C. *P. fluorescens* WCS365 wildtype or mutants alone are shown in black and co-inoculation with pathogens is shown in blue.

We tested protection against the N2C3 pathogen in two assays, one in hydroponics and one on the solid agar plates. Previous work identified some discrepancy between biofilm formation using these two assays, specifically that genes required for LPS modification were more important for colonization on solid surfaces [12, 26]. Similarly, in this study, we found that the solid rhizosphere condition altered the protection phenotype of certain biofilm mutants including those in the *wzt* and *clpP* genes. This finding might suggest that the function of these genes are more important for survival or competition with other microbes on solid surfaces or in an air-liquid interface assay.

We found that the mutants that failed to protect plants from the *P. fluorescens* N2C3 pathogen make similar, higher, or lower amounts of biofilm than wildtype *P. fluorescens* WCS365 on abiotic surfaces, indicating that the extent of *in vitro* biofilm biomass formation does not itself correlate with protection *in planta*. However, we observed a positive correlation between hypobiofilm formation *in vitro* and reduced plant weight, indicating that decreased biofilm production indeed diminishes the protective effect. Thus, the inability to properly form a biofilm *in vitro* impacts rhizosphere fitness of these mutants, which likely influences their capability to confer plant protection.

We found that loss of the large adhesion protein of *P. fluorescens* WCS365 (encoded by *lapDAEB* genes) resulted in strains unable to protect plants against *P. fluorescens* N2C3 and also formed significantly lower amounts of biofilm compared wildtype WCS365 (Figure 1D and Figure 3AB). The Lap system was previously shown to be required for biofilm formation in *P. fluorescens* WCS365 [21, 36]. The *lapA* gene encodes a large adhesion protein which is required for attachment to plant roots and abiotic surfaces [8, 9]. LapEB are components of the type I secretion system (T1SS) which is required for LapA secretion [16, 27]. The *lapD* gene, which encodes an inner membrane receptor of the intracellular signaling molecule c-di-GMP, determines the bacterial lifestyle transition between biofilm formation and dispersion by controlling the retention or the release of the LapA protein [16, 27]. Loss of protection by the *P. fluorescens* WCS365 *lapDAEB* mutants suggests that the entire Lap system and resulting secretion and cell-surface localization of LapA are likely necessary for protection against the *P. fluorescens* N2C3 pathogen, which may be due to the reduced rhizosphere colonization (Figure 2C).

Both biofilm formation and motility are important for successful rhizosphere colonization; however, they are often inversely regulated where downregulation of motility coincides with increase in biofilm forming ability [37–39]. As expected, the *fliK* (encodes a polar flagellar hook-length control protein) [20, 40], the *fliN* (encodes a flagellar motor switch protein) [41, 42], and the *fliP* (flagellar axial protein export apparatus) mutants [43] showed significantly increased biofilm formation (Figure 3A). Both *fliK* and *fliR* (flagellar axial protein export apparatus) mutants showed a loss of protection in the rhizosphere, suggesting that *P. fluorescens* WCS365 requires functional motility for rhizosphere colonization [34].

We found transposon insertions in a subset of lipopolysaccharides (LPS) modification genes lost protection against N2C3. As the major components of the Gram-negative bacterial outer membrane, lipopolysaccharides are more than just a physical permeability barrier for toxic compounds. LPS have been shown to be involved in versatile biological processes, such as host immunity recognition and evasion [44, 45], colonization [46, 47], and establishing symbiosis [48, 49]. LPS consists of three regions: the membrane anchoring region lipid A, the middle region core oligosaccharide, and the outside region referred to O-polysaccharide or O-antigen [50]. Genes involved in O-antigen transportation (*wzt*) [29] and biosynthesis (*wbpADE*, w*bpJ*) [51, 52] were identified in our mutant library. Wzt is part of the Wzm/Wzt ATP-binding cassette (ABC) transporter that specifically transports O serotypes O8 and O9a in *Escherichia coli* [29] and A-band O-antigen LPS in *Pseudomonas aeruginosa* PAO1[53]. The Wbp pathway including the *wbpADE* and w*bpJ* genes are in the B-band LPS serotype-specific O-antigen biosynthesis cluster in *P. aeruginosa* PAO1 [51–53]. While some components from the core oligosaccharide modification (*wapHR*), O-antigen transportation (*wzt*) [29] and biosynthesis (*wbpADE*, w*bpJ*) behaved differently under liquid or solid rhizosphere conditions, we found that the O-antigen of LPS plays a more important role in competition against the pathogen *in planta*.

Collectively, our work sheds light on the role of bacterial biofilm formation in colonization and plant pathogen protection in the rhizosphere. As *in vivo* biofilm is difficult to quantify, *in vitro* biofilm formation is often used as a proxy for *in vivo* biofilm production. Importantly, our screen of the *iam* and *sad* mutant libraries demonstrated that only a subset of genes involved in biofilm formation lost protection *in planta* indicating that *in vitro* biofilm formation does not always predict pathogen protection. Future characterization of genes that lost protection and have not previously been implicated in commensalism would also provide new insights into mechanisms of commensal-mediated protection of plants against pathogens. In addition, the characterization of the nonprotective mutants identified in this study will enhance our understanding of this process and provide better guidance to microbiome engineering.

## Materials and Methods

### Plant materials and growth conditions

*Arabidopsis thaliana* Col-0 seeds were surface sterilized by a mixture of 70% bleach followed by 10% ethanol, or 70% EtOH and 1.5% H_2_O_2_ solution on filter paper and allowed to dry for 30 min in a laminar flow hood. Sterilized seeds were then transferred into a centrifuge tube with 0.1% agar and stored in 4°C in the dark for 48 hrs before sowing. Seeds were germinated on square plates with ½ X Murashige and Skoog (MS) medium containing 1g/L 2-(N-morpholino) ethanesulfonic acid (MES) buffer, 2% sucrose, and 1% agar for 5 days. The pH of the MS medium was adjusted to 5.7 with 1M KOH. On day 6, plants of similar size were transferred to plates with ½ X MS medium containing 1 g/L MES buffer and 1% agar without sucrose. On day 7, plants were inoculated with 5 μL of bacterial culture with OD_600_ of 0.001. All the plant materials were grown at 22°C at 100 μM m^-2^s^-1^ light under a 12-hour light/dark cycle in a temperature-controlled growth room unless otherwise indicated.

### Bacterial strains and growth conditions

The library of *Pseudomonas fluorescens* WCS365 increased attachment mutants (*iam*) and surface attachment defective (*sad*) transposon insertion mutants were kindly provided by Dr. George O’Toole [14], from the Geisel School of Medicine at Dartmouth, USA. The library contains 65 mutants in total (Table S1). Overnight cultures were made in LB broth supplied with either 10 μg/mL gentamycin or 40 μg/mL tetracycline and grown at 28°C with shaking at 180 rpm. The bacterial overnight cultures were diluted and resuspended in 10 mM MgSO_4_ to the indicated OD_600_ prior to plant inoculation. *Pseudomonas fuscovaginae* SE-1 [54] was grown in LB at 28°C and *Pseudomonas aeruginosa* PAO1 [55] was grown in LB at 37°C.

### In vitro bacterial growth

The wildtype *P. fluorescens* WCS365 and the *iam* and *sad* libraries were grown overnight in LB broth in a 96-well plate. Optical density at 600 nm (OD_600_) was measured in a Biotek Epoch 2 plate reader. Cultures were diluted to an OD_600_ of 0.02 in fresh LB broth in 200 µL total in a 96 well plate. The plate was incubated and read in a Biotek Epoch 2 plate reader set to 28°C with continuous orbital shake for 24 hours. OD_600_ was read every 15 mins.

### Transposon insertion mapping by arbitrary PCR

The DNA sequences flanking the insertions in the transposon insertion mutants were determined by arbitrary PCR and Sanger sequencing as previously described [14, 56]. The DNA flanking regions were amplified by two rounds of PCR by using two sets of primers. In the first round of PCR, a primer Tn5Ext, which is unique to the transposon but more distal from the transposon end and an arbitrary primer (ARB1), which can hybridize the chromosomal sequences flanking the transposon were used to enrich the genomic DNA near the transposon. The second round of PCR used the PCR products from the first round of PCR as a template, a primer Tn5Int, which is also unique to the transposon but more proximal to the transposon end (around 60 bp from the transposon to the chromosome junction) and an ARB2 primer, which has identical 5’end as ARB1. The PCR products were purified either by PCR clean-up kit (QIAquick PCR purification kit) or by gel extract purification kit (QIAquick gel extract kit) and were then sent for Sanger sequencing. The insertion location was identified by BLAST using the *P. fluorescens* WCS365 reference genome (NCBI Accession CP089973.1).

### MYCroplanters

GFP-expressing *Pseudomonas fluorescens* N2C3 was generated using pSMC21 [57] via electropoartion. Breifly, an overnight culture of N2C3 in LB broth was pelleted and washed twice in 300 mM sucrose to generate electrocompetent cells. Transformed cells were plated on LB supplemented with 25 µg/mL kanamycin for selection.

Arabidopsis seeds were sterilized and germinated in the MYCroplanter system [25]. Five days after gemination, wildtype *P. fluorescens* WCS365 and the *iam* and *sad* libraries was grown overnight in LB broth in a 96-well plate at 30°C with shaking at 200 rpm. *P. fluorescens* N2C3-GFP was grown overnight in LB broth supplemented with 25 µg/mL kanamycin. On day 6, the WCS365 strains were spun down, spent LB broth was removed, and cells were resuspended in the same volume of ½ MS supplemented with MES and no sucrose. The N2C3-GFP culture was spun down, washed once to remove kanamycin, and resuspended in ½ MS supplemented with MES and no sucrose. For all strains, OD_600_ was measured using Biotek Epoch 2 plate reader. For transferring seedlings, a fresh 96-well plate was filled with 275 µL at a 1:1 ratio of wildtype WCS365 or *iam* or *sad* mutants and N2C3-GFP in 50,000 cells total using the cell estimate that 1 mL of *Pseudomonas* culture at OD_600_ of 1 is 5x10^8^ cells. A single MYCroplanter was transferred to each well of the 96-well plate, the lid brace was added, and the 96-well plate was sealed with 3M Micropore tape. The MYCroplanter system was incubator in a reach in growth chamber for 7 days. At harvest, MYCroplanters were transferred to the scanning tray and scanned on an Epson Perfection V850 Pro scanner as stated in [25]. GFP fluorescence and OD_600_ of the 96-well plate without MYCroplanters were read in a BioTek Synergy H1 plate reader. Health score calculations were performed as described in [25]. N2C3-GFP quantification was taken directly from the GFP fluorescence measurements. To calculate the fraction of WCS365 in the well, N2C3-GFP signal was converted to OD_600_ using a standard curve. For each well, the N2C3-GFP approximated OD_600_ was subtracted from the total OD_600_ reading to estimate the amount of WCS365 in the well.

### Pathogen protection assay on solid agar

The bacterial overnight cultures were diluted and resuspended in 10 mM MgSO_4_ to a final OD_600_ of 0.001 inoculum for single-strain inoculations. For bacterial competitions, bacterial mixtures were prepared in a ratio of 1:1 with final OD_600_ of 0.002 (0.001 for each strain). Each 7-day old plant was inoculated with 5 μL of bacterial culture along the root. Plates were dried in a biosafety cabinet before sealing and moved to the growth chamber as described above. Seven days later, all the plates were imaged using an Epson V850 flatbed scanner and weighed by pooling three plants per treatment.

### Crystal violet assay

Bacterial biofilm formation on a 96-well U-shape plastic surface was measured by a crystal violet assay [30]. Bacterial overnight cultures (as described above) were spun down at 10,000 × g for 3 min and the pellet was washed and resuspended in 1 × M63 medium. 100 μL of bacterial culture with OD_600_ of 0.1 was added in a 96-well plate with 8 technical replicates per treatment. Plates were subsequently incubated at 28°C for 18 hrs. After incubation, the bacterial culture was removed by inverting. The plate was then washed twice by gently submerging in a small tub of water and then inverting the plate to remove the water. 125 μL of a 0.1% crystal violet solution was added to the wells (including 8 wells that were used as background controls) and left for 10 min at room temperature. The plates were rinsed 3 times by submerging in water as described above. All residual water was removed by firmly shaking the plate and allowing it to dry at room temperature. For biofilm quantification, 125 μL of 30% acetic acid was added to each well including the blank wells to solubilize the crystal violet. After a 10 min incubation, 100 μL of the solubilized crystal violet or the 30% acetic acid was transferred to a new dish with flat bottom and the absorbance was read at 550 nm.

## Acknowledgments

This work was supported by NSERC Discovery Grant (NSERC-RGPIN-2021-03587) and CIHR Grant (PJT - 169051) to C.H.H and NIH grant (R01-GM123609) to G.A.O. Additional trainee support was provided by a Chinese Scholarship Council Award to Y.L., and a CGS-M award to Z.Y., and an NSERC Banting Postdoctoral fellowship to M.Y.C.

---

**Figure S1.**
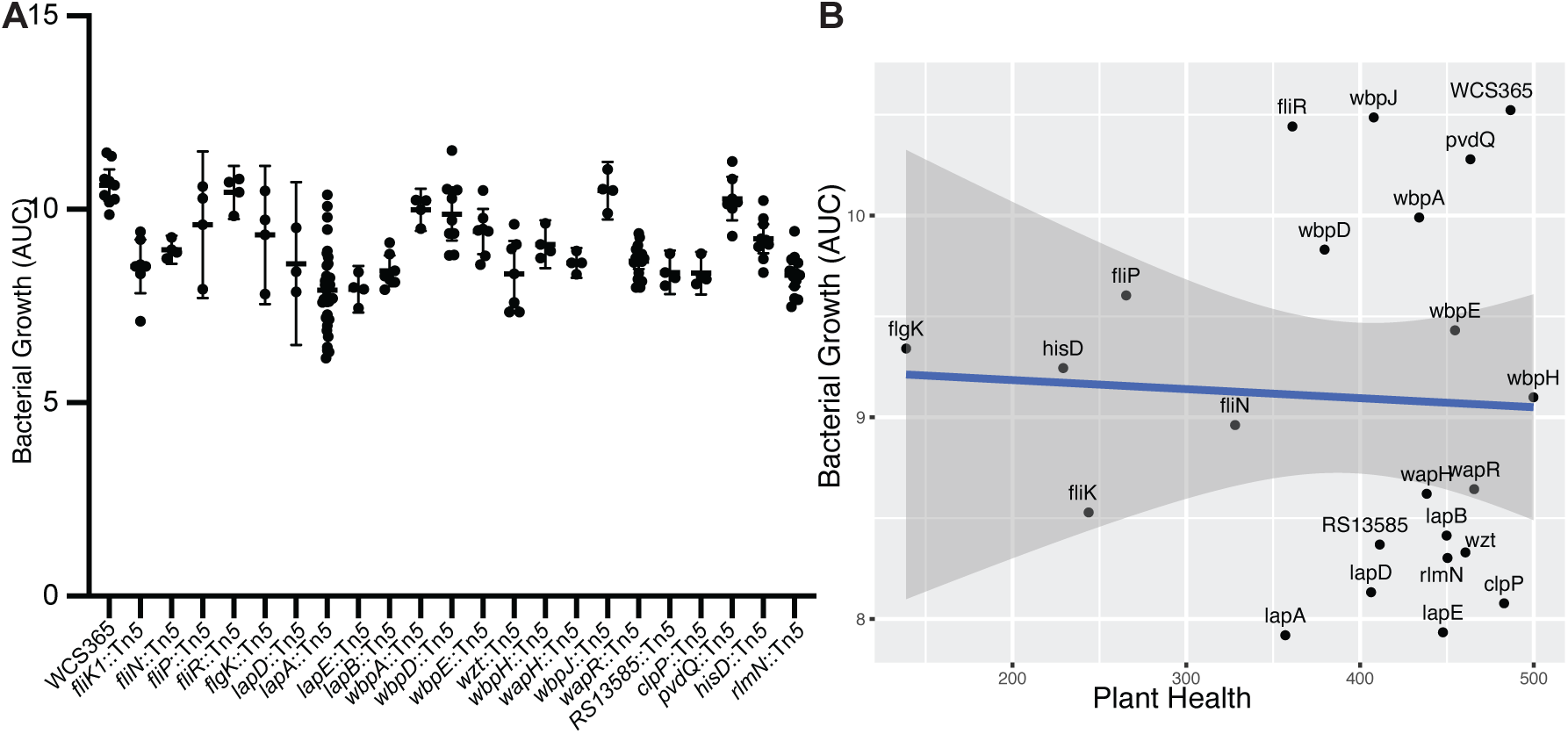
*In vitro* and *in planta* growth of mutants. A) Transposon insertion mutants were grown in vitro in LB medium and the area under the curve was quantified. B) A linear regression was performed between the ability to protect plants from pathogens and *in vitro* bacterial growth and no significant correlation was noted.

**Table S1.** Insertion sites of transposon insertions in the *iam* and *sad* libraries.

| Original Mutant name | Well | Seq published | Gene name | Locus Tag | Gene position | Figure 3 |
| --- | --- | --- | --- | --- | --- | --- |
| *iam-1::Tn5Gent* | A1 | This study | *wapR1* | LRP86_RS13590 | 696(885) | Yes |
| *iam-2::Tn5Gent* | A2 | This study | *wapH* | LRP86_RS06880 | 2273(3243) | Yes |
| *iam-3::Tn5Gent* | A3 | This study | *wapR2* | LRP86_RS13590 | 676(885) | No |
| *iam-5::Tn5Gent* | A4 | This study | *wbpH* | LRP86_RS06855 | 449(1242) | No |
| *iam-6::Tn5Gent* | A5 | This study | *wapR3* | LRP86_RS13590 | 697(885) | No |
| *iam-7::Tn5Gent* | A6 | This study | *wapR4* | LRP86_RS13590 | 688(885) | No |
| *iam-8::Tn5Gent* | A7 | This study | *LRP86_RS13585* | LRP86_RS13585 | 667(1142) | Yes |
| *iam-9::Tn5Gent* | A8 | This study | *wapR5* | LRP86_RS13590 | 697(824) | No |
| *iam-10::Tn5Gent* | A9 | This study | *wapR6* | LRP86_RS13590 | 676(885) | No |
| *iam-11::Tn5Gent* | A10 | This study | *wbpJ* | LRP86_RS06850 | 216(1227) | Yes |
| *sad-7::Tn5Gent* | C1 | This study | *lapE* | LRP86_RS12055 | 987(1350) | Yes |
| *sad-8::Tn5Gent* | C2 | This study | *lapA* | LRP86_RS12045 | 994(14239) | No |
| *sad-9::Tn5Gent* | C3 | This study | *lapA* | LRP86_RS12045 | 2755(14239) | No |
| *sad-10::Tn5B30(Tc)* | C4 | This study | *HP1* | LRP86_RS10235 | 1323(1415) | No |
| *sad-11::Tn5B30(Tc)* | C5 | O'Toole 1998 | *clpP1* | LRP86_RS20465 | 29(636) | Yes |
| *sad-12::Tn5B30(Tc)* | C6 | This study | *proA* | LRP86_RS07155 | 82(1272) | No |
| *sad-13::Tn5B30(Tc)* | C7 | O'Toole 1998 | *fliP* | LRP86_RS18760 | 403(750) | Yes |
| *sad-14::Tn5B30(Tc)* | C8 | O'Toole 1998 | *flgK* | LRP86_RS18635 | 255(2016) | Yes |
| *sad-15::Tn5B30(Tc)* | C9 | This study | *pvdQ* | LRP86_RS25805 | 385(2349) | Yes |
| *sad-16::Tn5B30(Tc)* | C10 | This study | *fliK* | LRP86_RS18735 | 307(1374) | Yes |
| *sad-17::Tn5B30(Tc)* | C11 | This study | *fliK* | LRP86_RS18735 | 371(1374) | No |
| *sad-18::Tn5B30(Tc)* | C12 | O'Toole | *lapA* | LRP86_RS12045 | 6411(14239) | No |
| *sad-19::Tn5B30(Tc)* | D1 | O'Toole 2006 | *lapD* | LRP86_RS12025 | 1470(1947) | Yes |
| *sad-20::Tn5B30(Tc)* | D2 | This study | *fliN* | LRP86_RS18750 | 299(459) | Yes |
| *sad-21::Tn5B30(Tc)* | D3 | This study | *pvdQ* | LRP86_RS25805 | 369(2349) | No |
| *sad-22::Tn5B30(Tc)* | D4 | This study | *HP2* | LRP86_RS23120 | -54(498) | No |
| *iam-21::Tn5B30(Tc)* | B1 | This study | *wbpE* | LRP86_RS06900 | 372(1092) | Yes |
| *iam-21::Tn5B30(Tc)* | A11 | This study | *wbpE* | LRP86_RS06900 | 372(1092) | No |
| *iam-22::Tn5B30(Tc)* | B2 | This study | *wzt* | LRP86_RS06885 | 204(1785) | No |
| *iam-22::Tn5B30(Tc)* | A12 | This study | *wzt* | LRP86_RS06885 | 204(1785) | Yes |
| *iam-23::Tn5B30(Tc)* | B3 | This study | *wbpD* | LRP86_RS06905 | 508(585) | Yes |
| *iam-24::Tn5B30(Tc)* | B4 | This study | *wbpA* | LRP86_RS06915 | 592(1309) | Yes |
| *iam-25::Tn5B30(Tc)* | B7 | This study | *wbpD* | LRP86_RS06905 | 508(585) | No |
| *iam-26::Tn5B30(Tc)* | B8 | This study | *wbpD* | LRP86_RS06905 | 508(585) | No |
| *sad-43::Tn5B30(Tc)* | D5 | This study | *HP3* | no hits |  | No |
| *sad-44::Tn5B30(Tc)* | D6 | This study | *HP4* | LRP86_RS23825 | 314(627) | No |
| *sad-45::Tn5B30(Tc)* | D7 | This study | *bioF* | LRP86_RS08345 | 137(1179) | No |
| *sad-46::Tn5B30(Tc)* | D8 | This study | *mexW* | LRP86_RS03915 | 1709(3027) | No |
| *sad-47::Tn5B30(Tc)* | D9 | This study | *lapA4* | LRP86_RS12045 | 12307(14239) | No |
| *sad-48::Tn5-B30 (Tc)* | D10 | This study | *fliR* | LRP86_RS18770 | 544(783) | Yes |
| *sad-51::Tn5-B22 (Gm, 'lacZ)* | D11 | This study | *lapA5* | LRP86_RS12045 | 4385(14239) | No |
| *sad-52::Tn5-B22 (Gm, 'lacZ)* | D12 | This study | *lapB* | LRP86_RS12060 | 1095(2157) | Yes |
| *sad-53* | E11 | This study | *lapA6* | LRP86_RS12045 | 4136(14239) | No |
| *sad-55::Tn5-B22 (Gm, 'lacZ)* | C2 | This study | *rlmN1* | LRP86_RS15910 | 1106(1149) | No |
| *sad-56::Tn5-B22 (Gm, 'lacZ)* | C3 | This study | *rlmN2* | LRP86_RS15910 | 956(1149) | No |
| *sad-57::Tn5-B22 (Gm, 'lacZ)* | C4 | This study | *rlmN3* | LRP86_RS15910 | 1113(1149) | No |
| *sad-58::Tn5-B22 (Gm, 'lacZ)* | E5 | This study | *morA* | LRP86_RS06630 | 4245(4248) | No |
| *sad-62::Tn5-B22 (Gm, 'lacZ)* | E6 | This study | *lapA7* | LRP86_RS12045 | 4136(14239) | No |
| *sad-63::Tn5-B22 (Gm, 'lacZ)* | E7 | This study | *HP5* | LRP86_RS05960 | 1273(1725) | No |
| *sad-79::Tn5-B22 (Gm, 'lacZ)* | E8 | This study | *lapA8* | LRP86_RS12045 | 11171(14139) | Yes |
| *sad-80::Tn5-B22 (Gm, 'lacZ)* | E9 | This study | *lapA9* | LRP86_RS12045 | 11261(14239) | No |
| *sad-81::Tn5-B22 (Gm, 'lacZ)* | E10 | This study | *lapA10* | LRP86_RS12045 | 11179(14239) | No |
| *sad-82::Tn5-B22 (Gm, 'lacZ)* | E11 | This study | *lapA11* | LRP86_RS12045 | 11182(14239) | No |
| *sad-83::Tn5-B22 (Gm, 'lacZ)* | E12 | This study | *rpfC* | LRP86_RS23115 | 828(1638) | Yes |
| *sad-84::Tn5-B22 (Gm, 'lacZ)* | F1 | This study | *lapE* | LRP86_RS12055 | 1241(1350) | No |
| *sad-86::Tn5-B22 (Gm, 'lacZ)* | F2 | This study | *lapA12* | LRP86_RS12045 | 4386(14239) | No |
| *sad-87::Tn5-B22 (Gm, 'lacZ)* | F3 | This study | *lapB* | LRP86_RS12060 | 1744(2157) | No |
| *sad-89::Tn5-B22 (Gm, 'lacZ)* | F4 | This study | *hisD* | LRP86_RS15545 | 443(1338) | No |
| *sad-95::Tn5-B22 (Gm, 'lacZ)* | F5 | This study | *hisD* | LRP86_RS15545 | 442(1338) | No |
| *sad-96::Tn5-B22 (Gm, 'lacZ)* | F6 | This study | *hisD* | LRP86_RS15545 | 442(1338) | No |
| *sad-97::Tn5-B22 (Gm, 'lacZ)* | F7 | This study | *moaA* | LRP86_RS02125 | 799(969) | No |
| *sad-98::Tn5-B22 (Gm, 'lacZ)* | F8 | This study | *proA* | LRP86_RS07155 | 264(1272) | No |
| *sad-100::Tn5-B22 (Gm, 'lacZ)* | F9 | This study | *rlmN4* | LRP86_RS15910 | 911(1149) | No |
| *sad-101::Tn5-B22 (Gm, 'lacZ)* | F10 | This study | *sadC* | LRP86_RS04535 | 184(1107) | Yes |
| *sad-102::Tn5-B22 (Gm, 'lacZ)* | F11 | This study | *metF* | LRP86_RS08870 | intergenic | No |

## Notes

### Competing Interest Statement

The authors have declared no competing interest.

